# A Novel Method for Calculating Mean Erythrocyte Age Using Erythrocyte Creatine

**DOI:** 10.1101/642942

**Authors:** Masashi Kameyama, Masafumi Koga, Toshika Okumiya

## Abstract

**Backgrounds:** Erythrocyte creatine (EC) decreases reflecting erythrocyte age.

**Methods:** We developed an EC model, which showed a bi- or mono-exponential relationship between mean erythrocyte age (*M*_*RBC*_) and *EC*. We reanalyzed the previously published data of 21 patients with hemolytic anemia which included *EC* and ^51^Cr half-life.

**Results:** *M*_*RBC*_ and log_*e*_ *EC* showed excellent significant linearity (*r* = −0.9475, *p <* 0.001), showing that it can be treated as a mono-exponential relationship within the studied range (*EC*: 1.45 – 11.76 *µ*mol/g Hb). We established an equation to obtain *M*_*RBC*_ (days) from *EC* (*µ*mol/g Hb), *M*_*RBC*_ = −22.84 log_*e*_ *EC* + 65.83.

**Conclusion:** This equation allows calculation of *M*_*RBC*_ based on EC which has practical applications such as the diagnosis of anemia.

## Introduction

Estimating the lifespan of erythrocytes is useful for the differential diagnosis of anemia as it is known that the erythrocyte lifespan of hemolytic patients is shortened [1]. Previously, obtaining the lifespan or mean age of erythrocytes was very difficult, therefore it was seldom measured. Furthermore, supply of ^51^Cr, which is needed for measuring erythrocyte lifespan, was ceased in Japan in 2015, due to low demand. This left Japanese doctors unable to measure the erythrocyte lifespan of patients by means of ^51^Cr. Biotin-label [2, 3] is also used to measure the erythrocyte lifespan, however its procedure is very troublesome as well, requiring aseptic labeling of erythrocyte and repeated blood samplings. Breath carbon monoxide (CO) measurement [4, 5] may be useful to estimate erythrocyte turnover; however, this technique cannot be applied to smokers. We proposed a method to estimate erythrocyte mean age from HbA1c and average glucose [6]. However the method needs a glycation constant to be determined by another method. Some indices such as reticulocyte and haptogloblin does not seem to be sensitive to indicate mild hemolysis. Cases with latent hemolysis were reported who showed normal reticulocyte, normal haptogloblin and yet shortened erythrocyte life-span [7–9].

Creatine in the cells is maintained by creatine transporters. Deficiency in this transporter leads to symptoms [10, 11]. Young erythrocytes have adequate transporter activity resulting in intracellular creatine being tens of times higher than in plasma. However, the activity of the transporter gradually diminishes, so that old erythrocytes cannot maintain this concentration gradient.

Erythrocyte creatine (EC) has been demonstrated to be an excellent indicator of hemolysis [12, 13]. Mean erythrocyte age (*M*_*RBC*_) estimation using EC would be more convenient than the ^51^Cr method as it requires only one blood sample. However, though an increase in EC value has been correlated with shorter lifespan of erythrocytes, EC value itself has not previously been used for the estimation of *M*_*RBC*_ directly. An estimation of *M*_*RBC*_ would be more useful for quantitative assessment of patients than simple EC value. Moreover, *M*_*RBC*_ derived by EC may be comparable with *M*_*RBC*_ derived by other methods.

Here, we would like to propose an equation to obtain *M*_*RBC*_ from EC concentration based on a model.

## Methods

### Patients

Data from 21 patients with hemolytic anemia that was published by Fehr *et al.* [13] was used. As this is a re-analysis study, approval by the institutional review board was not required.

### Data conversion

We estimated *M*_*RBC*_ by multiplying half life of ^51^Cr by 2.61. As human erythrocytes do not obey the Poisson process [14], the term “half-life” is not entirely suitable for erythrocytes. Fehr *et al.* [13] determined ^51^Cr half-life, the elution-corrected ^51^Cr half-life, and the mean cell lifespan. The mean cell lifespan was not recorded in their table. The elution-corrected ^51^Cr half-life would provide an estimate of *M*_*RBC*_, considering that normal erythrocytes in a human have a similar lifespan [14]. However, their elution-corrected ^51^Cr half-life seems less concordant with EC rank. Complicated procedure sometimes reduce the stability of the system. Therefore, we chose the simple uncorrected ^51^Cr half-life in the same way as Fehr *et al.* [13]. Considering that *M*_*RBC*_ for normal erythrocytes is about 60 days, and their normal range of ^51^Cr half-life was 23 – 27 days, multiplying ^51^Cr half life by 2.61 (= 60*/*23) provides a good estimation of *M*_*RBC*_ in practice.

The units for erythrocyte creatine concentration used in Fehr *et al.* [13], mg/dL of red cells was converted into *µ*mol/g Hb by the following equation, assuming mean cell hemoglobin concentration (MCHC) is 33g/dL. The molecular weight of creatine is 131.15. While MCHC varies naturally and also reduces in iron deficiency anemia, variability in MCHC is generally low.

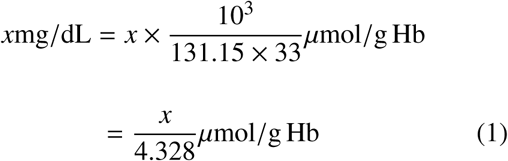

### Data analysis

Data on *EC* and *M*_*RBC*_ were analyzed with spreadsheet software, Excel^®^. 365 (Microsoft Corporation, Redmond, WA, USA).

Logarithms of *EC* and *M*_*RBC*_ was plotted considering our model (Supplement).

## Results

### Relatiohship between *M*_*RBC*_ and log_*e*_ *EC*

A significant linear relationship (*r* = −0.9475, df =19, *t* = 12.92, *p* = 7.368 × 10^-11^) was observed between ^51^Cr-derived *M*_*RBC*_ and log_*e*_ *EC* (Figure 1). The relationship appears to be mono-exponential which is concurrent with the prediction by model (*See* Supplement) that the relationship would be bi- or mono-exponential.

**Figure 1.**
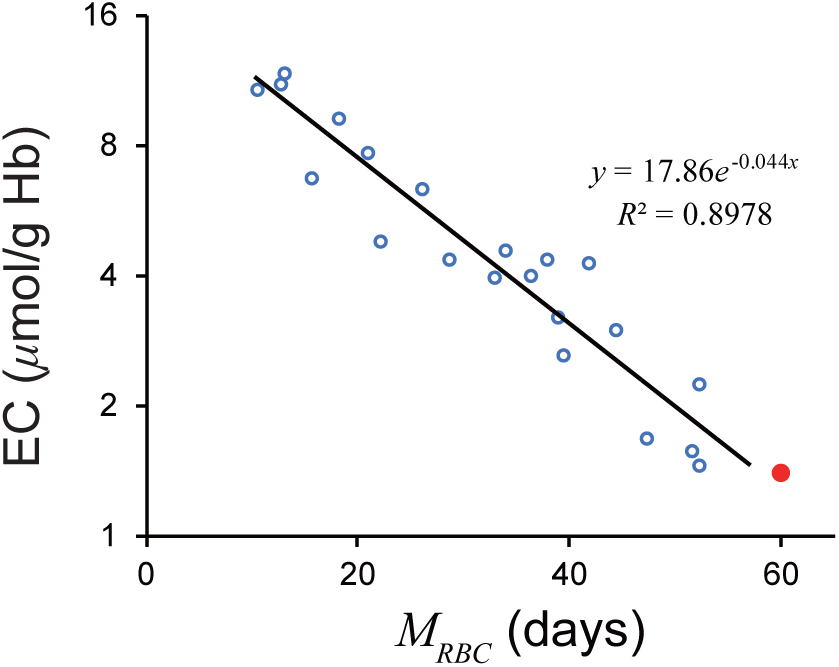
Relatiohship between *M*_*RBC*_ and log*_e_ EC*. A significant linear relationship was observed. A *red closed circle* denotes a standard value; *M*_*RBC*_ = 60 days, *EC* = 1.4*µ*mol/g Hb. A *black line* denotes a regression line. *EC*, erythrocyte creatine; *M*_*RBC*_, mean erythrocyte age.

A regression line was as follows.

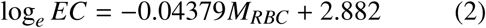

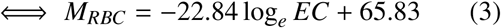

A standard value of *EC*, 1.4 *µ*mol/g Hb gives an *M*_*RBC*_ of 58.14 days.

Equation (3) accurately estimated *M*_*RBC*_ from *EC* values (Figure 2)

**Figure 2.**
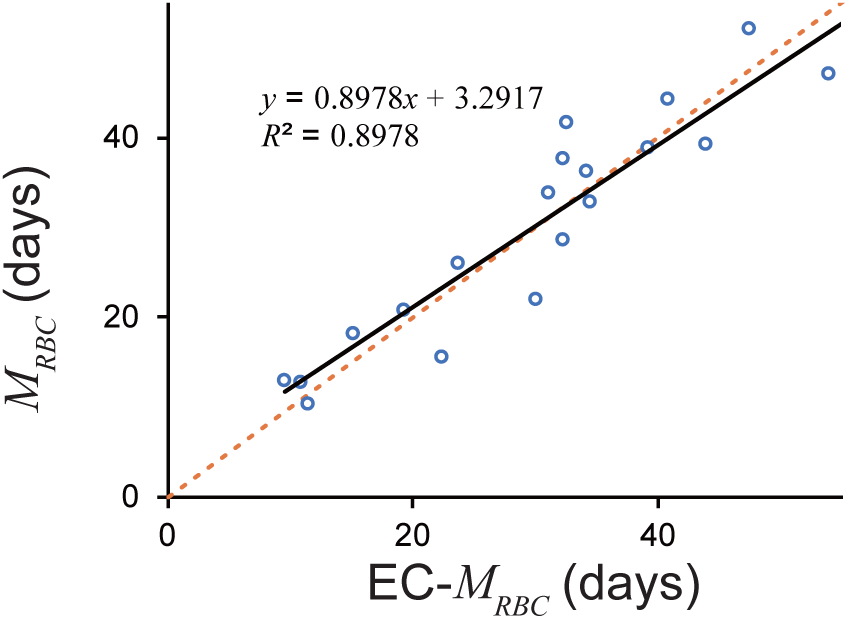
*M*_*RBC*_ estimated by *EC* and 51Cr. EC derived *M*_*RBC*_ showed excellent estimation. An *orange dotted line* denotes line of identification (*y* = *x*). A *black line* denotes a regression line. *EC*, erythrocyte creatine; *M*_*RBC*_, mean erythrocyte age.

## Discussion

The current study successfully established a reliable method of estimating *M*_*RBC*_ from *EC* based on a creatine model (Supplement). We would be able to determine a glycation constant for the method to estimate erythrocyte mean age from HbA1c and average glucose.

Although Fehr *et al.* [13] divided patients into a severe hemolytic disease group and a group with milder forms of hemolysis, our model revealed that logarithm of *EC* can combine the two groups (Figure 3). The regression formula passed close to a standard value of *EC*, 1.4 *µ*mol/g Hb and 60 days of *M*_*RBC*_, which proves the validity of the formula.

**Figure 3.**
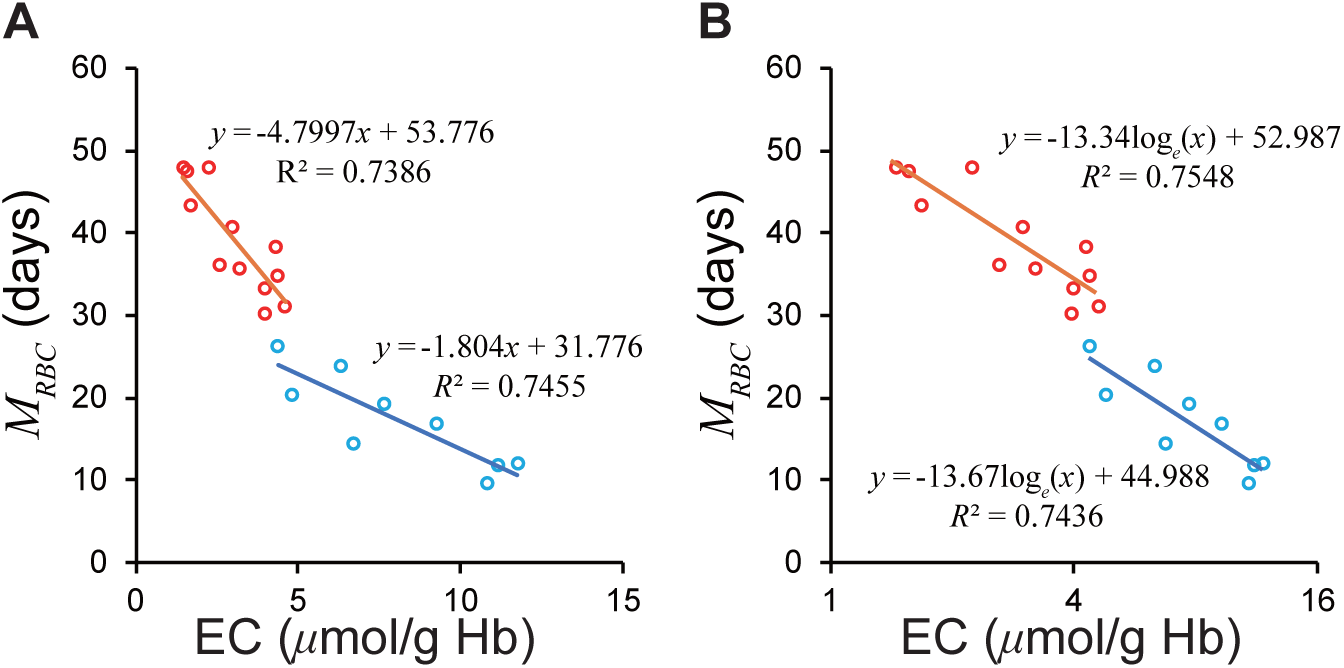
Relationship between *EC* and *M*_*RBC*_ in the groups with severe and mild hemolytic disease. (A) The two groups show differing regression lines on a normal scale. (B) The two groups are unified on a semi-logarithmic scale. The *Red circles* represent mild group, *sky blue* the severe group according to Fehr *et al.* [13]. *EC*, erythrocyte creatine; *M*_*RBC*_, mean erythrocyte age.

It cannot be determined which wing of the two lines (Supplement) the observed line of the log_*e*_ *EC* – *M*_*RBC*_ relationship is on; *i.e.* whether the slope of the graph represents the rate constant for creatine diffusion (*λ*_1_) or the rate constant for decline in creatine transporter (*λ*_2_). Another equation may need to be developed for value ranges not explored in this study.

The devised method was formulated entirely based on the previously presented data from 21 patients. This method should be verified by further study with various hematological diseases including thalassemia and hereditary spherocytosis. Estimation of *M*_*RBC*_ from ^51^Cr half-life may not be optimal, although we believe that it would be tolerable. The EC transporter activity function, 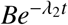 relies solely on the assumption that the number of transporters reduces over-time randomly due to erythrocytes’ lack of nucleus. However, the linear relationship between log_*e*_ *EC* and *M*_*RBC*_ confirms the assumption. The EC measuring method of Fehr *et al.* [13] used a diacetyl-*l*-naphthol chemical reaction, which is less sensitive than the recently developed *N*-methylcarbamoyl derivative of methylene blue, 10-*N*-methylcarbamoyl-3,7-*bis*(dimethylamino)phenothiazine (MCDP) enzyme method [15]. Further study on the validity of our proposed formula would be best done in a country where ^51^Cr is available.

## Conclusion

Our equation allows calculation of *M*_*RBC*_ based on EC which has practical applications such as the diagnosis of anemia.

## Acknowledgments

This study is partly supported by Grants-in-Aid for Scientific Research, 18K07488 for MKameyama. The authors would like to thank Ms. Natalie Okawa for English language editing of this manuscript.

