## Supplement for "A Novel Method for Calculating Mean Erythrocyte Age Using Erythrocyte Creatine"

### Theory

The terms in the equations are summarized in Table 1.

#### Creatine in a single erythrocyte

The rate of diffusion of creatine would be proportional to the concentration of creatine in the cell. We assume that the transporter activity obeys an exponential function; *i.e.* transporters diminish randomly (proportional to the number of the transporter) and are not renewed due to lack of nucleus.

$$\frac{dCr}{dt} = -\lambda_1 Cr(t) + Be^{-\lambda_2 t} \tag{1}$$

where $Cr(t)$ denotes creatine concentration in a $t$-day-old erythrocyte, $\lambda_1$ denotes rate constant of creatine diffusion. $Be^{-\lambda_2 t}$ ($B > 0$) is creatine transporter activity. This differential equation can be solved analytically if $\lambda_1 \neq \lambda_2$.

$$\frac{d(Cr - \frac{B}{\lambda_1-\lambda_2}e^{-\lambda_2 t})}{dt} = -\lambda_1 Cr(t) + \frac{B\lambda_1}{\lambda_1 - \lambda_2}e^{-\lambda_2 t} \tag{2}$$

$$Cr(t) = Ae^{-\lambda_1 t} + \frac{B}{\lambda_1 - \lambda_2}e^{-\lambda_2 t} \tag{3}$$

where $A$ denotes an integral constant.

$Cr(t)$ is a sum of two exponential functions, it can be treated as a bi-exponential function or a mono-exponential function within a range of interest.

The fate of $Cr(t)$ is dependent on whether $\lambda_1 - \lambda_2 > 0$ or not. If $\lambda_1 - \lambda_2 < 0$ and $C'(0) > 0$, $Cr(t)$ has a peak when $t > 0$ (Figure 1 *orange line*). It can be treated as a mono-exponential function after the second term is negligible. However, as EC monotonically and rapidly decreases after birth of the erythrocytes [1], we can exclude this condition.

If $\lambda_1 - \lambda_2 < 0$ and $C'(0) \leq 0$, $Cr(t)$ decreases monotonically. As $\lambda_2 > \lambda_1$, $e^{-\lambda_2 t}$ decreases more rapidly. Moreover, the second negative term must be small, considering that $C'(0) < 0$ yields $\frac{B}{\lambda_2-\lambda_1} < \frac{\lambda_1}{\lambda_2}A$. Therefore, the second negative term can be considered negligible comparing the first term.

If $\lambda_1 - \lambda_2 > 0$, $Cr(t)$ is a bi-exponential function. The logarithm of a bi-exponential function can be expressed by a bent line (Figure 1B *blue line*), because a large $t$ makes one term negligible, while small $t$ ($\to -\infty$) makes the other term negligible.

*, Department of Diagnostic Radiology, Tokyo Metropolitan Geriatric Hospital and Institute of Gerontology, 35-2 Sakae-cho, Itabashi-ku, Tokyo, 173-0015, Japan
†Department of Internal Medicine, Hakuhokai Central Hospital, Amagasaki, 661-0953, Japan
‡Department of Biomedical Laboratory Sciences, Faculty of Health Sciences, Kumamoto University, Kumamoto, 860-8556, Japan

Table 1: Terms used in the text

| Term | Definition | Representative value |
|---|---|---|
| $Cr(t)$ | creatine concentration in a $t$-day-old erythrocyte | |
| $\lambda_1$ | rate constant for creatine diffusion | |
| $\lambda_2$ | rate constant for decline in creatine transporter | |
| $\lambda$ | substitute for $\lambda_1$ or $\lambda_2$ | |
| $A, B, C, D$ | constants | |
| $EC$ | mean erythrocyte creatine concentration | 1.4 $\mu$mol/g Hb |
| $\alpha$ | a parameter of gamma distribution | 25.59 |
| $\beta$ | a parameter of gamma distribution | 5.59 |
| $p(t)$ | probability density function of RBC death | |
| $R_0$ | erythrocyte production rate | – /day |
| $R(t)$ | the number of erythrocytes at $t$ days after birth | |
| $M_{RBC}$ | mean red blood cell age | 60 days |
| $RBC$ | number of erythrocytes | – |

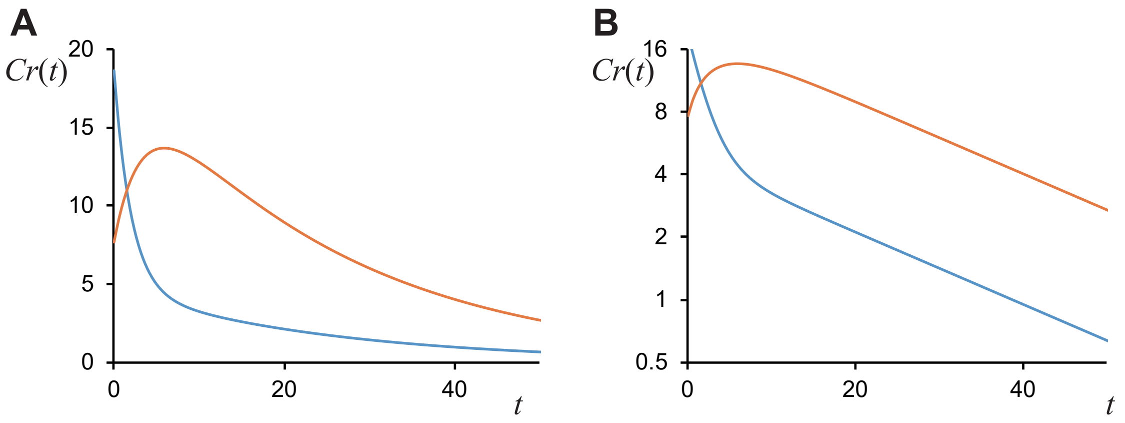


Figure 1: Examples of $Cr(t)$. (A) normal scale; (B) logarithmic scale. *Orange line:* $\lambda_1 - \lambda_2 < 0$ and $C'(0) > 0$, the function has a peak. *Blue line:* $\lambda_1 - \lambda_2 > 0$, $Cr(t)$ is a bi-exponential function. The logarithm of a bi-exponential function can be expressed by a bent line.

#### Problem applying single cell model to erythrocyte population

Equation (3) itself would not be suitable to obtain mean erythrocyte age, because EC is not measured from a single cell.

$$EC = \left(\sum_i^n Cr(t_i)\right)/n$$

$$= A\left(\sum_i^n e^{-\lambda_1 t_i}\right)/n + \frac{B}{\lambda_1 - \lambda_2}\left(\sum_i^n e^{-\lambda_2 t_i}\right)/n \qquad (4)$$

$y = e^{-\lambda x}$ is downward convex. The centroid of $n$-polygonal, $(t_i, e^{-\lambda t_i})$ is over the curve of $y = e^{-\lambda x}$.

$$\left(\sum_i^n e^{-\lambda t_i}\right)/n \geq \exp\left(-\lambda \frac{\sum_i^n t_i}{n}\right) \qquad (5)$$

Therefore, when $\lambda_1 - \lambda_2 > 0$,

$$EC \geq Ae^{-\lambda_1 M_{RBC}} + \frac{B}{\lambda_1 - \lambda_2} e^{-\lambda_2 M_{RBC}} \qquad (6)$$

#### Erythrocyte lifespan

Kameyama *et al.* [2] have recently calculated RBC lifespan based on the probability density function $p(t)$ of RBC death proposed by Shrestha *et al.* [3].

$$p(t) = \frac{1}{\Gamma(\alpha)\beta^{\alpha}} t^{\alpha-1} e^{-t/\beta} \tag{7}$$

$\Gamma$ denotes the Euler gamma function.

$$\Gamma(\alpha) = \int_0^{\infty} x^{\alpha-1} e^{-x} dx \tag{8}$$

The number of erythrocytes ($RBC$) and mean erythrocyte age ($M_{RBC}$) was calculated. (*See* Kameyama *et al.* [2] for details.)

$$RBC = R_0 \int_0^{\infty} tp(t)dt = R_0 \alpha\beta \tag{9}$$

$$M_{RBC} = \frac{(\alpha+1)\beta}{2} \tag{10}$$

#### Creatine model

The number of $t$-day-old erythrocyte is $R(t)$. Each RBC has $Ae^{-\lambda_1 t} + \frac{B}{\lambda_1-\lambda_2} e^{-\lambda_2 t}$ creatine. Therefore, mean creatine concentration, $EC$ can be described as follows:

$$EC = \int_0^{\infty} R(t) \times \left( Ae^{-\lambda_1 t} + \frac{B}{\lambda_1 - \lambda_2} e^{-\lambda_2 t} \right) dt / RBC \tag{11}$$

$$\int_0^{\infty} R(t) \times e^{-\lambda t} dt = \left[ R(t) \frac{e^{-\lambda t}}{-\lambda} \right]_0^{\infty} - \int_0^{\infty} R'(t) \frac{e^{-\lambda t}}{-\lambda} dt = \frac{R_0}{\lambda} - \frac{R_0}{\lambda} \int_0^{\infty} p(t) e^{-\lambda t} dt \tag{12}$$

$$\int_0^{\infty} p(t) e^{-\lambda t} dt = \frac{1}{\Gamma(\alpha)\beta^{\alpha}} \int_0^{\infty} t^{\alpha-1} e^{-(1/\beta+\lambda)t} dt = \frac{1}{\Gamma(\alpha)\beta^{\alpha}} \frac{\Gamma(\alpha)}{(1/\beta+\lambda)^{\alpha}} = \frac{1}{(1+\beta\lambda)^{\alpha}} \tag{13}$$

Hence, $EC$ can be expressed as follows:

$$EC = \frac{A}{\lambda_1 \alpha\beta} \left( 1 - \frac{1}{(1+\beta\lambda_1)^{\alpha}} \right) + \frac{B}{\lambda_1 - \lambda_2} \frac{1}{\lambda_2 \alpha\beta} \left( 1 - \frac{1}{(1+\beta\lambda_2)^{\alpha}} \right) \tag{14}$$

#### Approximation of the derived relationship

The Taylor expansion provides the following equation:

$$(1+\beta\lambda)^{-\alpha} \simeq 1 - \alpha\beta\lambda + \frac{\alpha(\alpha+1)}{2} (\beta\lambda)^2 - \frac{\alpha(\alpha+1)(\alpha+2)}{6} (\beta\lambda)^3 + \ldots \tag{15}$$

$$\frac{1}{\lambda\alpha\beta} \left( 1 - \frac{1}{(1+\beta\lambda)^{\alpha}} \right) \simeq \frac{1}{\lambda\alpha\beta} \left( \alpha\beta\lambda - \frac{\alpha(\alpha+1)}{2} (\beta\lambda)^2 + \frac{\alpha(\alpha+1)(\alpha+2)}{6} (\beta\lambda)^3 - \ldots \right) = 1 - \frac{\beta(\alpha+1)}{2} \lambda + \frac{(\alpha+1)(\alpha+2)}{6} (\beta\lambda)^2 - \ldots \tag{16}$$

As $\alpha \gg 1$,

$$\frac{1}{\lambda\alpha\beta} \left( 1 - \frac{1}{(1+\beta\lambda)^{\alpha}} \right) \simeq 1 - \lambda M_{RBC} + \frac{2}{3} \lambda^2 M_{RBC}^2 - \ldots \tag{17}$$

Thus, $\frac{1}{\lambda\alpha\beta} \left( 1 - \frac{1}{(1+\beta\lambda)^{\alpha}} \right)$ can be described approximately as a function of $M_{RBC}$. Although how $\alpha$ and $\beta$ vary when $M_{RBC}$ decreases or increases cannot be determined, this implies that the function $\frac{1}{\lambda\alpha\beta} \left( 1 - \frac{1}{(1+\beta\lambda)^{\alpha}} \right)$ would not be greatly affected by $\alpha$ and $\beta$ if $M_{RBC} = (\alpha + 1)\beta/2$ is satisfied. This can be confirmed numerically (Figure 2A). Calculations based on the assumption that

$\beta$ was constant and $\beta$ was proportionate to $\alpha$ showed similar results. Therefore, $\beta$ can be considered a constant.

Below, $\frac{1-e^{-x}}{x}$ was approximated to be $De^{-Cx}$, as the shape of the graphs were similar (Figure 2B).

$$C = \frac{-x_0 e^{-x_0} + (1 - e^{-x_0})}{x_0(1 - e^{-x_0})}, \qquad D = \frac{1 - e^{-x_0}}{x_0 e^{-Cx_0}} \quad (18)$$

provides the same value of the two function and the differential function at $x = x_0$. Figure 2B visualizes the approximation.

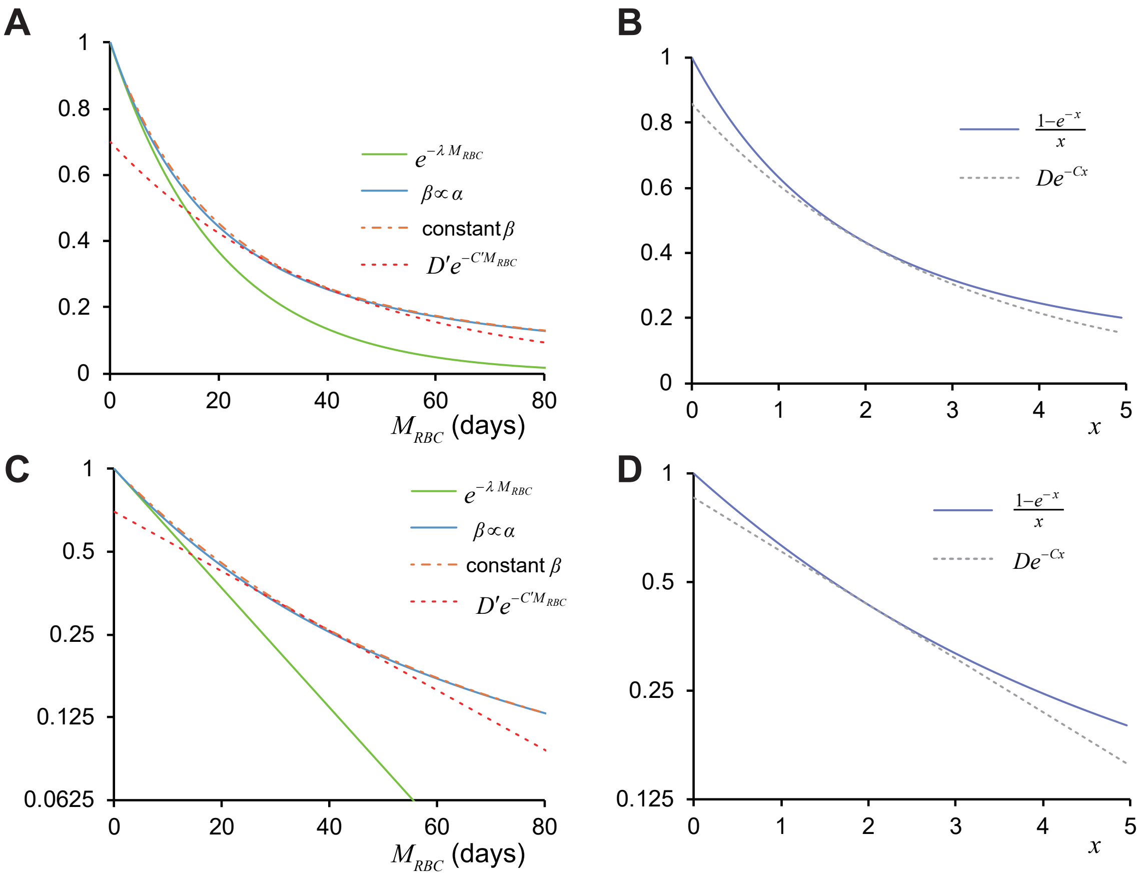


Figure 2: (A) Relatiohship between $M_{RBC}$ and $\frac{1}{\lambda\alpha\beta}\left(1 - \frac{1}{(1+\beta\lambda)^\alpha}\right)$. The two condition of $\frac{1}{\lambda\alpha\beta}\left(1 - \frac{1}{(1+\beta\lambda)^\alpha}\right)$ (constant $\beta$ and $\beta \propto \alpha$) showed similar results. Note that $\frac{1}{\lambda\alpha\beta}\left(1 - \frac{1}{(1+\beta\lambda)^\alpha}\right)$ is consistently larger than $e^{-\lambda M_{RBC}}$. $D'e^{-C'M_{RBC}}$ is an approximation at $M_{RBC} = 36.33$ with exponential function by equation (19). (B) $\frac{1-e^{-x}}{x}$ and its approximation, $De^{-Cx}$ (equation (18)) when $x_0 = 2$. (C, D) Semi-log scales of (A, B) show that $\frac{1-e^{-x}}{x}$ and $\frac{1}{\lambda\alpha\beta}\left(1 - \frac{1}{(1+\beta\lambda)^\alpha}\right)$ can be treated as an exponential function.

As $\frac{1-e^{-x}}{x}$ can be treated as an exponential function, it follows that $\frac{1}{\lambda\beta x}\left(1 - \frac{1}{(1+\beta\lambda)^x}\right)$ can also be treated as an exponential function, because

$$\frac{1}{\lambda\beta\alpha}\left(1 - \frac{1}{(1+\beta\lambda)^\alpha}\right) = \frac{\log_e(1+\beta\lambda)}{\lambda\beta}\frac{1 - \exp(-\log_e(1+\beta\lambda)\alpha)}{\log_e(1+\beta\lambda)\alpha}$$

$$\simeq \frac{\log_e(1+\beta\lambda)}{\lambda\beta} D \exp\left(-C \log_e(1+\beta\lambda)\left(\frac{2M_{RBC}}{\beta}-1\right)\right) \tag{19}$$

$C, D$ can be estimated by equation (18). To obtain an approximation around $M_{RBC0}$, $x_0$ should be as following:

$$x_0 = \log_e(1+\beta\lambda)\left(\frac{2M_{RBC0}}{\beta}-1\right) \tag{20}$$

In conclusion, $EC$ can be expressed approximately as a (bi-)exponential function of $M_{RBC}$.
